# Directed fMRI-based Functional Connectivity Estimation using Physics-Informed Neural Networks

**DOI:** 10.1101/2024.07.09.602748

**Authors:** Roberto C. Sotero, Jose M. Sanchez-Bornot

## Abstract

Estimating directed functional connectivity (dFC) within the brain is crucial for comprehending neural interactions. However, conventional methodologies encounter constraints in accuracy, scalability, and interpretation. The method presented here harnesses Physics-Informed Neural Networks (PINNs) to amalgamate the governing physical principles of brain dynamics, thereby improving dFC estimation from resting-state functional magnetic resonance imaging (rsfMRI) data. In particular, during the training phase, we derive the input weights from a long-short term memory (LSTM) network, which, within our framework, represent the influence of all other brain areas on the specific region under consideration. These input weights are then integrated into the nonlinear differential equation that models the rsfMRI time series within the specific brain area. Through the training of the PINN model, we simultaneously estimate, for each brain area, the biophysical parameters of the model, including the dFC parameters from all the remaining areas. We applied this methodology to both autism spectrum disorder (ASD) and neurotypical data, revealing significant sex-specific differences in connectivity patterns. These findings underscore the potential of PINNs in advancing our understanding of neural dynamics and emphasize the significance of directionality in brain connectivity research.

## 1 INTRODUCTION

The exploration of brain function through the lens of connectivity has become a central theme in neuroscience, offering important insights into the complex interplay of neural circuits in both health and disease [1]. Functional connectivity, a term coined to describe the statistical dependencies between different neurophysiological events, has emerged as a pivotal concept in understanding the brain’s intricate communication networks [2]. The utilization of functional connectivity measures has significantly advanced our comprehension of the brain’s organizational principles, elucidating the functional architecture that underpins cognitive processes and characterizes neuropsychiatric disorders [3], [4].

Traditionally, functional connectivity has been quantified using Pearson correlation (PC) [2], a method that captures the linear relationship between the temporal dynamics of neurophysiological events across different brain regions. This approach, while invaluable in highlighting the strength and significance of pairwise interactions, inherently lacks directional information, presenting a notable limitation in deciphering the flow of information within the brain’s networks. The absence of directionality in PC-based functional connectivity analyses restricts our ability to infer causal relationships between neural events, ultimately impacting our understanding of the brain’s functional hierarchy and the coordination of its distributed components.

The limitations of nondirectional functional connectivity measures have catalyzed the development and adoption of directed functional (dFC) and effective connectivity methods, designed to infer the direction and, potentially, the causality of neuronal interactions. Techniques such as Granger causality, dynamic causal modeling (DCM), and transfer entropy have been introduced to address the directional aspect of functional interactions, offering a window into the causal architecture of brain networks [5], [6], [7]. These methods provide insights into the directional flow of information, allowing researchers to construct more detailed models of neural communication and to better understand the regulatory mechanisms underlying cognitive functions and their dysregulation in diseases.

However, dFC analyses come with their own set of challenges and limitations. The computational complexity and the need for extensive time series data to achieve reliable estimates pose significant hurdles. Furthermore, the assumptions inherent in model-based approaches, such as DCM [6], may not always be valid across different contexts or states of brain activity, potentially leading to misinterpretations of the directional interactions. Thus, while functional connectivity analyses have undeniably enriched our understanding of the brain’s functional organization, the transition from nondirectional to directional measures introduces both opportunities and challenges. Enhancing the accuracy and interpretability of dFC measures remains a crucial frontier in neuroscience research, promising to deepen our understanding of the brain’s operational principles in health and disease.

In this paper, we introduce a novel approach to studying dFC through the integration within a Physics-Informed Neural Network (PINN). PINNs are an innovative category of machine learning models that assimilate physical laws, articulated through differential equations, into the framework of neural networks [8]. These networks undergo training that encompasses not only data-driven learning but also adherence to the foundational physical principles governing the system under investigation. The integration of these physical constraints ensures that PINN predictions are aligned with established physics, thereby improving their precision and applicability, even in situations characterized by limited or noisy data [8]. PINNs have emerged as powerful tools for solving both forward and inverse problems across various fields, including fluid dynamics, materials science [9], and, more recently, neuroscience [10]. They offer a promising framework for modeling complex biological processes that are based on well-established physical principles [11]. In this context, we have adapted a Brain Dynamics Model (BDM) for the Blood Oxygen Level-Dependent (BOLD) signal [10] observed in resting-state functional magnetic resonance imaging (rsfMRI) studies. Our approach involves integrating the neural network’s input weights and adjusting these weights using a sign matrix that is derived from the PC matrix, in addition to a region-specific parameter. This technique allows for the precise mapping of information flow direction and enables the concurrent estimation of dFC to a targeted brain region from all other regions. Such large-scale dFC estimation is beyond the capabilities of previous biophysical models, highlighting the unique advantage of the proposed methodology.

## 2 MATERIALS AND METHODS

### 2.1 Data preprocessing and preparation

In this study, we analyzed rsfMRI data from the Autism Brain Imaging Data Exchange (ABIDE-I) dataset registered in 17 international sites [12]. The dataset comprises 1112 subjects, of which 539 have ASD (474 males and 65 females), and 573 are neurotypical (474 males and 99 females). Detailed demographics information is presented in [13]. To ensure consistency and avoid the variability introduced through different preprocessing pipelines, we selected datasets preprocessed with the Configurable Pipeline for the Analysis of Connectomes (CPAC), as provided by ABIDE. The details for this preprocessing procedure are provided in the literature [14].

We employed different preprocessing methods for rsfMRI data to enhance the accuracy of ASD classification, in line with recommendations from prior studies [15], [16]. Our preprocessing pipeline incorporated bandpass filtering within 0.01 to 0.1 Hz and Global Signal Regression (GSR) to minimize non-neural noise. To account for the variance in BOLD-rsfMRI data stemming we standardized the signals by resampling each participant’s brain area time series to a two-second temporal resolution and segmenting them into two-minute epochs. We then applied Z-score normalization to each BOLD time series to achieve standardization.

For each participant, the PC matrix was calculated from the time series data of 116 brain regions delineated by the Automated Anatomical Labeling (AAL) atlas [17]. To determine the statistical significance of the functional connections, we applied the False Discovery Rate (FDR) correction [18], setting the threshold for significance at an FDR-adjusted p-value of 0.05. Connections that did not meet this criterion were considered non-significant and set to zero. Consequently, we derived the sign matrix *S* by taking the sign of the non-zero PC coefficients.

### 2.2 LSTM networks

Long Short-Term Memory (LSTM) networks are a type of Recurrent Neural Network (RNN) designed to recognize patterns in sequences of data [19]. The core concept behind an LSTM network is the cell state, which acts as a kind of conveyor belt to carry relevant information throughout the processing of the sequence. LSTM units include a set of gates that regulate the flow of information into and out of the cell state. The equations that govern the behavior of the LSTM unit at each time step *t* are:

- Forget Gate:

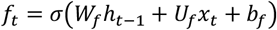
- Input Gate:

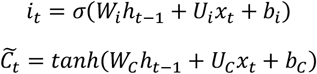
- Cell State Update:

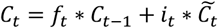
- Output Gate:

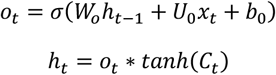

where *W* are the weights associated with the input *x*_*t*_ for each of the gates and the cell state, *U* are the weights associated with the recurrent connections, *b* represents the bias term for each gate, *σ* is the sigmoid function and *tanh* is the hyperbolic tangent. *f*_*t*_, *i*_*t*_, *o*_*t*_, and 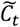 are the outputs of the forget gate, input gate, output gate, and the candidate cell state, respectively. *C*_*t*_ and *C*_*t*−1_ are the current and previous cell states, respectively. *h*_*t*_ and *h*_*t*−1_ are the current and previous hidden states, respectively. The asterisk (*) denotes element-wise multiplication.

### 2.3 Brain Dynamics Model of the rsFMRI signal

We begin with a recently developed BDM [10], which characterizes the time series of the i-th brain area, represented by *y*_*i*_(*t*), using the equation:

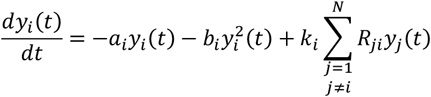

Here the −*a*_*i*_*y*_*i*_ (*t*) with *a*_*i*_ > 0 indicates linear negative feedback while the nonlinear term 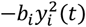 counterbalances the grow of *y*_*i*_ (*t*). The summation 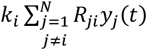 represents the cumulative effect of all other brain regions on the i-th region, modulated by the functional connectivity matrix *R*_*ji*_, which is computed from the data. *N* is the number of brain areas. The factor *k*_*i*_ can be interpreted as the degree of receptiveness of region *i* to inputs from other regions.

### 2.4 dFC estimation within the PINN framework

In this paper, we present a new framework for estimating dFC by integrating it within a PINN. This approach capitalizes on the strengths of PINNs, which is the inclusion of physical laws—here, the differential equations that model BOLD signals—into the training process of the neural network. The resulting model excels not only in predicting the dynamics of BOLD signals but also in simultaneously estimating the intricate connectivity patterns and biophysical parameters that govern these dynamics.

Central to our model are LSTM networks, selected for their proficiency in capturing time series fluctuations, making them especially suitable for handling BOLD-rsfMRI signals. The neural network architecture we use here encompasses two LSTM layers, with a dropout layer interposed to mitigate overfitting. Following the final LSTM layer is a fully connected layer tasked with predicting the BOLD signal for a specific brain area *i*, denoted as *y*_*i*_ (*t*), at each time point *t*. The number of LSTM hidden units is the same in the two layers and is denoted as *N*_*u*_; the dropout rate is *p*. The inputs to the network consist of a matrix that combines the time vector with the time series from all brain areas, excluding area *i*, and has dimensions *N* × *T*, where *T* is the length of the BOLD time series. The LSTM weights, *W*_*f*_,*W*_*i*_, *W*_*C*_, and *W*_*o*_, are fundamental to the network’s input processing. During the training process, we derive the vector *w* as the amalgamation of the absolute values of the input weight matrices, averaged over the LSTM units:

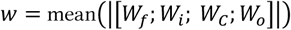

The first row, corresponding to the temporal feature, is discarded to preserve only the influences from the *N* − 1 brain areas. The influence of each area *j* on area *i* is represented by *w*_*ji*_, obtained from *w*, and is used to calculate *dFC*_*ji*_ as follows:

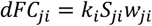

where *S*_*ji*_ is the sign matrix derived from the PC Matrix. Then, the BDM that constrains the PINN is written as:

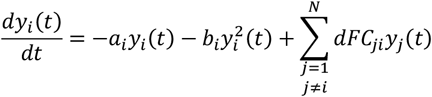

Within this framework (Figure. 1), we simultaneously estimate parameters *a*_*i*_, *b*_*i*_, and the *N* − 1 values of *dFC*_*ji*_. The neural network’s performance is assessed using a Mean Square Error (MSE) loss function, which integrates two distinct components: Data Loss and Physics Loss. Data Loss evaluates the discrepancy between the predicted and observed BOLD signals, while Physics Loss ensures the predictions adhere to the dynamics defined by the brain dynamics model’s differential equation. The network’s goal is to estimate parameters that fit the observed data while complying with the established physical laws. The PINN was implemented using PyTorch [20] and trained for a maximum of 5000 epochs. To mitigate overfitting, we utilized dropout layers along with *L*2 regularization, and instituted early stopping if the validation loss failed to improve after 100 epochs. The Adam optimizer [21] is used for training the network, with the learning rate (*α*) being treated as a hyperparameter. Finally, Bayesian optimization through the ‘hyperopt’ library [22] is used to fine-tune the model’s four hyperparameters: the number of hidden units, the dropout rate, the learning rate, and the *L*2 regularization coefficient.

**Figure 1:**
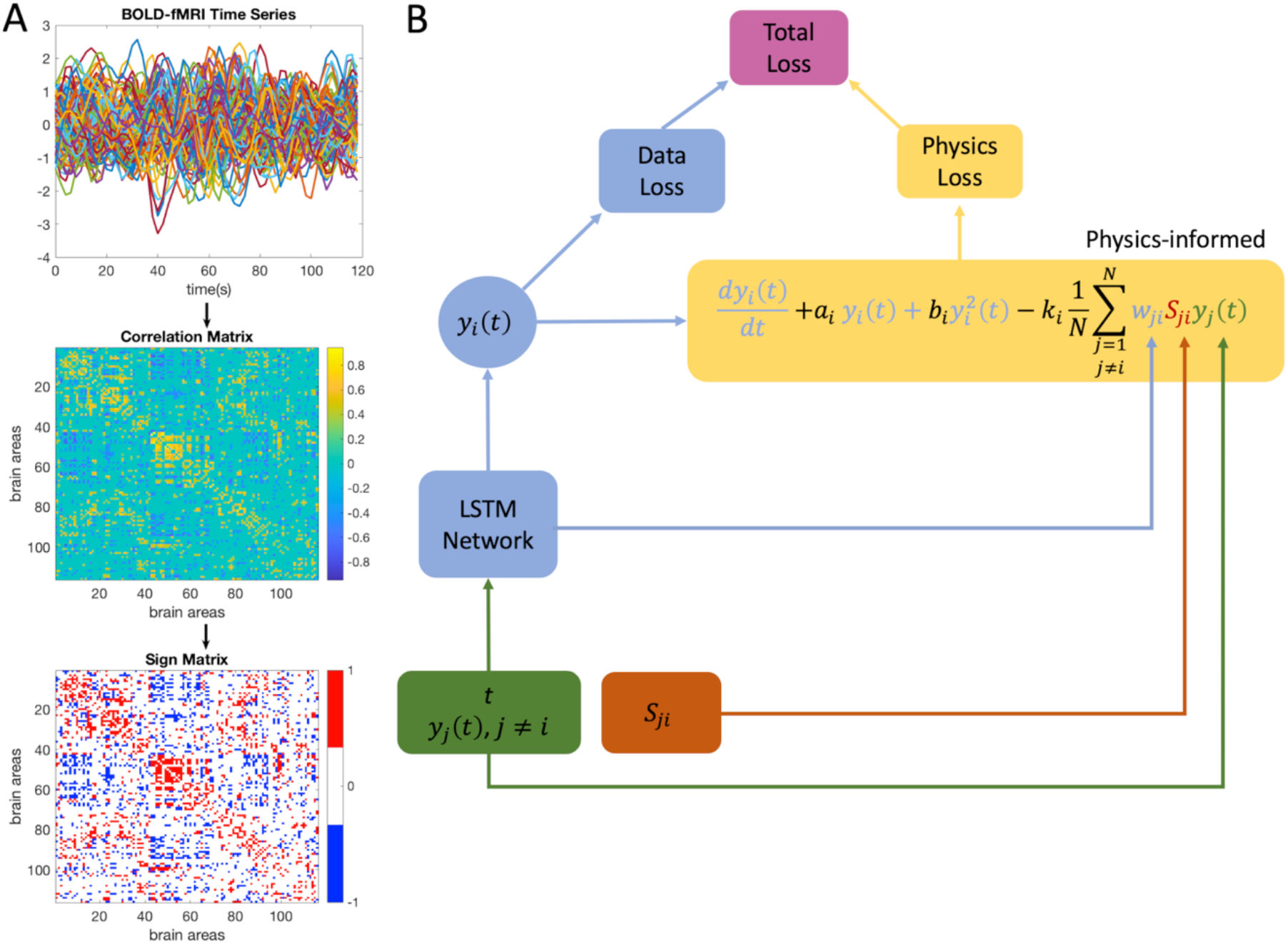
Framework of the Physics-Informed Neural Network (PINN) for estimating directed Functional Connectivity (dFC). A) Preprocessing of BOLD-fMRI Data: The process begins with the normalized BOLD-fMRI time series *y*_*i*_ (*t*) from 116 brain regions of an individual subject. We compute the Pearson Correlation Matrix *R*_*ji*_, which leads to the formation of the Sign Matrix *S*_*ji*_. B) Architecture of the Model: the time vector *(t*) and the time series of all regions (*y*_*j*_(*t*), *j* ≠ *i*), are the inputs to the LSTM network that predicts *y*_*i*_(*t*) for each specific brain region. The Total Loss function is a composite measure consisting of Data Loss and Physics Loss. Data Loss evaluates the discrepancy between the predicted BOLD signals and the observed fMRI data. Physics Loss ensures compliance with the underlying dynamical systems modeled by the differential equation. This equation integrates both linear and nonlinear BOLD signal dependencies with coefficients *a*_*i*_ and *b*_*i*_, respectively. It also includes a term accounting for the mean influence from other regions, parameterized by *k*_*i*_, and the learned input weight coefficients *w*_*ji*_ from the LSTM network, modulated by the Sign Matrix *S*_*ji*_. The dFC values are then computed as *dFC*_*ji*_ = *k*_*i*_*S*_*ji*_*w*_*ji*_.

## 3 RESULTS

Figure. 2 reveals notable distinctions in the estimated parameters *a*_*i*_ and *b*_*i*_ for certain brain regions when comparing individuals with ASD to neurotypical (NT) controls. In the male cohort (row A), a comparative analysis (neurotypical minus autism) of the parameter *a*_*i*_—representative of the linear rate of change in the BOLD signal—unveils significant variances in only seven out of 116 regions examined. Regions such as the right Frontal Superior Orbital, left Postcentral, right Supramarginal, and right Angular exhibit a more marked linear dynamic in the neurotypical group. Conversely, in the right Amygdala, left Cerebellum 9, and Vermis 8, this linear dynamic is more accentuated in the autism group. For the parameter *b*_*i*_, which signifies non-linear dynamics, positive mean differences are observed in eight regions, with the most pronounced disparities in the right Frontal Superior Orbital, left Postcentral, and right Supramarginal areas, while negative mean differences are noted in three regions, including the left Cerebellum 9, right Cerebellum 8, and left Lingual.

**Figure 2:**
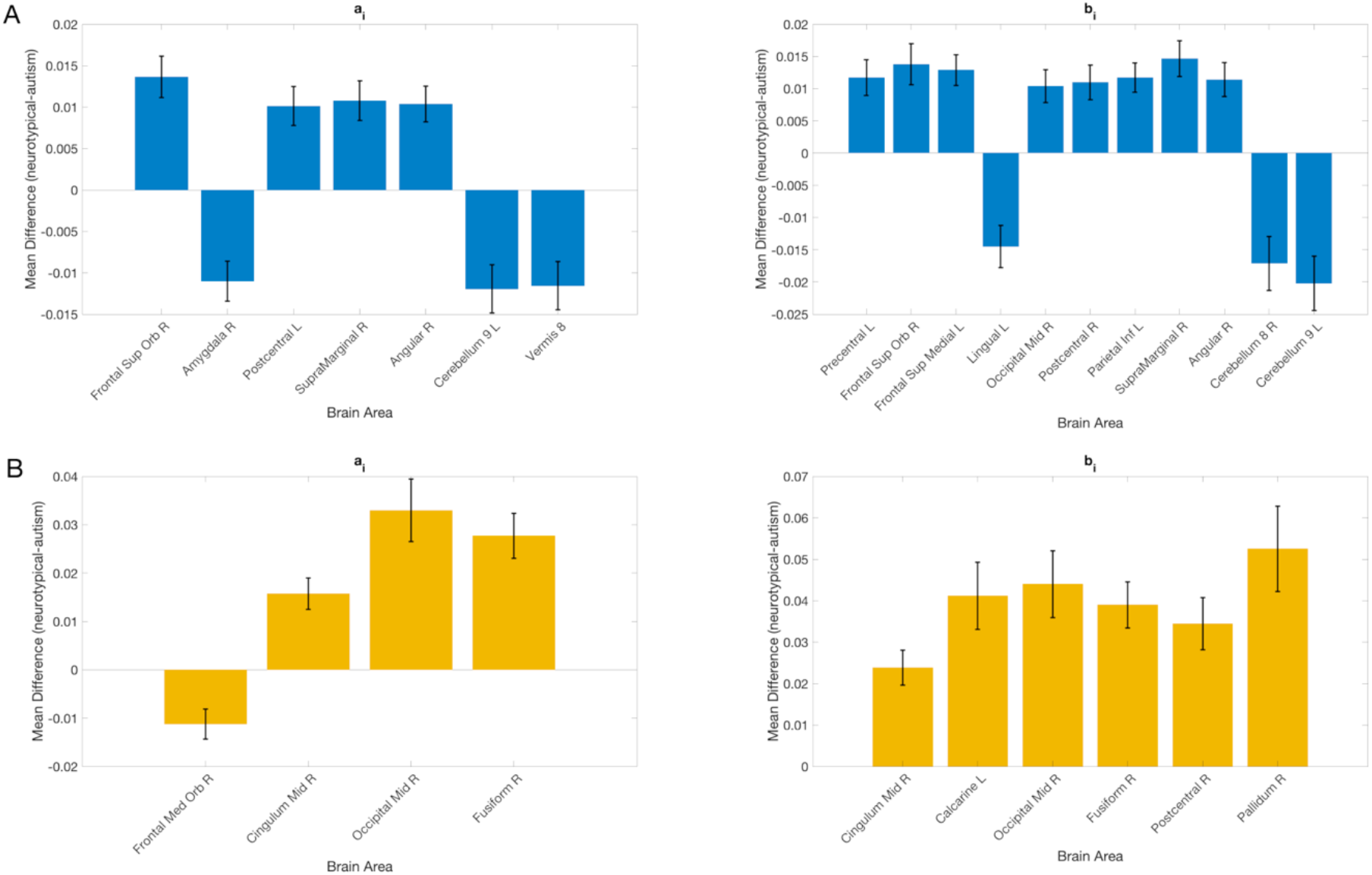
Sex-specific parameter estimation from rsfMRI data using PINNs. A) Male cohort: The bar graphs illustrate the significant mean differences in the estimated parameters *a*_*i*_, *b*_*i*_ and *k*_*i*_ across brain regions for the male group, comparing individuals with ASD to neurotypical controls. Error bars indicate the standard error of these estimates, underscoring the variability within each group. B) Female cohort.

In the female cohort (row B), four regions exhibit significant differences for the linear parameter *a*_*i*_, with the right Frontal Medial Orbital showing negative differences, and the right Occipital Middle, right Fusiform, and right Cingulate Middle presenting positive differences. As for the non-linear parameter *b*_*i*_, all significant differences are positive, and they include the right Cingulate Middle, left Calcarine, right Occipital Middle, right Fusiform, right Postcentral, and right Pallidum. These findings indicate distinct linear and non-linear dynamics between male and female cohorts. However, it is important to acknowledge the varying ASD prevalence between genders, which is mirrored in the ABIDE dataset’s disproportionately higher number of male subjects compared to females. The specific brain regions demonstrating significant differences are particularly noteworthy, aligning with previous research that has pinpointed many of these areas as critical in the context of autism. For example, the Amygdala, known for its role in social and emotional processing, has been frequently associated with the pathophysiology of autism [23]. Similarly, the Cerebellum’s involvement in motor functions and cognitive processes has been a focal point in studies examining the developmental trajectory of autism [24]. The Frontal Superior Orbital region, which showed pronounced linear dynamic differences in our study, has been linked to executive function and behavior regulation, both of which can be atypical in autism [25]. Additionally, the Supramarginal and Angular gyri, associated with language and cognitive functions, may also contribute to the communication difficulties experienced by individuals with autism [26], [27]. Furthermore, structural and functional abnormalities in the Fusiform gyrus, often referred to as the ‘face area’, have been correlated with challenges in face recognition and social interaction in autism [28].

Figure. 3 displays the contrast in connectivity strengths between dFC and PC across seven resting-state networks [29]: Default Mode (DMN), Salience (SAL), Frontoparietal (FP), Sensory (SENS), Visual (VIS), Cerebellar (CEREB), and Temporal-Brainstem-Globus Pallidus (TBG). The analyses are conducted over four distinct subgroups: ASD males, ASD females, NT males, and NT females. Panel A focuses on the positive values within the connectivity matrices, showcasing modest differences (dFC-PC) for both ASD and NT groups across genders, with the FP and SENS networks demonstrating slightly larger disparities. Panel B considers the negative values within the connectivity matrices, where more pronounced dFC-PC differences emerge, particularly in the VIS, SENS, and SAL networks. This suggests that the conventional PC approach may underestimate the strength of negative connections compared to dFC. Panel C shows the absolute values of all connections in the connectivity matrices, revealing a similar pattern to Panel B, emphasizing that the applied dFC method predominantly modifies the negative connections relative to the PC matrix. When comparing subgroups, NT females and NT males exhibit the most significant differences between dFC and PC within the VIS network, whereas the largest discrepancies for the SENS network occur in both ASD male and female subgroups. Notably, for both positive and negative connections, the SENS network consistently shows among the largest differences. In summary, the results from Fig. 2 and Figure. 3 suggest sex-specific distinctions in connectivity alterations in autism, in agreement with existing literature on sex differences in ASD prevalence and neural architecture.

**Figure 3:**
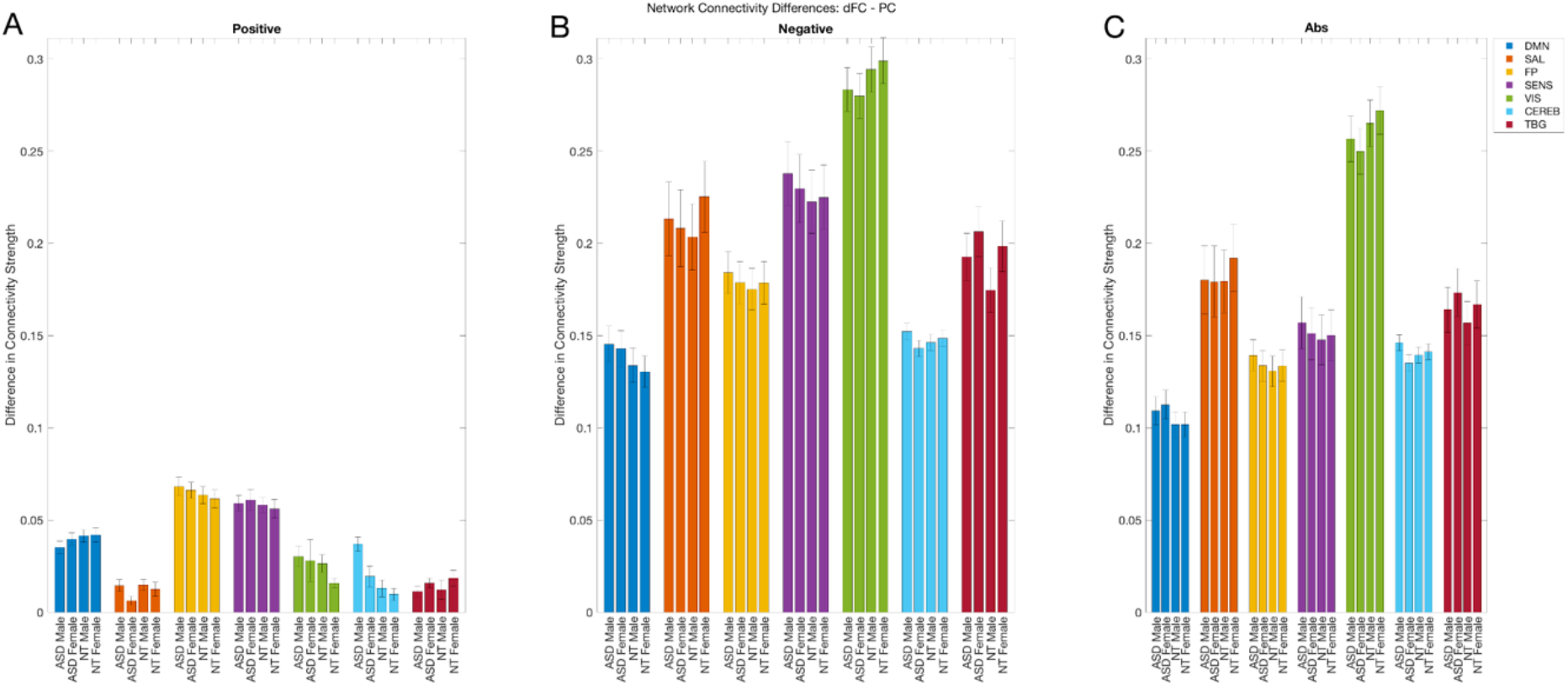
Network-Specific dFC-PC Differences by Condition. Analysis of connectivity variations within seven distinct neural networks, categorized by condition (ASD Male, ASD Female, Neurotypical Male, Neurotypical Female). Subplots A, B, and C focus on the quantification of positive, negative, and the absolute value of all connections, respectively, comparing dynamic functional connectivity (dFC) to Pearson correlation (PC) metrics. Each bar, color-coded for network identification, represents the mean difference in connectivity for each condition across the networks: Default Mode Network (DMN), Salience (SAL), Frontoparietal (FP), Sensory (SENS), Visual (VIS), Cerebellar (CEREB), and Temporal-Brainstem-Globus Pallidus (TBG). Error bars indicate the standard error of these estimates.

## 4 CONCLUSIONS

In this study, we presented a novel approach utilizing physics-informed neural networks for estimating directed functional connectivity within the brain. Our model overcomes traditional biophysical modeling limitations by concurrently estimating all connections to a particular brain region. This improvement significantly broadens the model’s adaptability for complex neural systems, thus facilitating enhanced insights into brain dynamics and potentially fostering the development of sophisticated diagnostic tools for neuroscience.

## ACKNOWLEDGMENTS

This work was supported by grant RGPIN-2022-03042 from Natural Sciences and Engineering Council of Canada. The authors are grateful for access to the Tier 2 High-Performance Computing resources provided by the Northern Ireland High Performance Computing (NI-HPC) facility funded by the Engineering and Physical Sciences Research Council (EPSRC), Grant No. EP/T022175/1.

## Notes

### Competing Interest Statement

The authors have declared no competing interest.

